# Fornix subdivisions and spatial learning: a diffusion MRI study

**DOI:** 10.1101/2025.05.15.654313

**Authors:** Carl J. Hodgetts, Mark Postans, Angharad N. Williams, Kim S. Graham, Andrew D. Lawrence

**Author notes:** Joint first authors (Carl J. Hodgetts, Mark Postans). Correspondence to Professor Andrew D. Lawrence.

## Abstract

The fornix is the major white matter tract linking the hippocampal formation with distal brain sites. Human and animal lesion studies show that the connections comprising the fornix are vital for specific attributes of episodic and spatial memory. The fornix, however, interconnects the hippocampal formation with an array of subcortical and cortical sites and it is not known which specific connections support spatial-mnemonic function. To address this, making use of a partly previously published dataset (Hodgetts et al., 2020), we applied a novel deterministic tractography protocol to diffusion-weighted magnetic resonance imaging (dMRI) data from a group of healthy young adult humans who separately completed a desktop-based virtual reality analogue of the Morris water maze task. The tractography protocol enabled the two main parts of the fornix, delineated previously in axonal tracing studies in rodents and primates, to be reconstructed in vivo, namely the pre-commissural fornix (connecting the hippocampus to the medial prefrontal cortex and the basal forebrain) and the post-commissural fornix (connecting the hippocampus to the medial diencephalon). We found that inter-individual differences in pre-commissural – but not, surprisingly, post-commissural – fornix microstructure (indexed by free water corrected fractional anisotropy, FA) were significantly correlated with individual differences in spatial learning, indexed by reduction in search error as individuals learned to navigate to a hidden target location from multiple starting points. This study provides novel evidence that flexible and/or precise spatial learning involves a hippocampal-basal forebrain/prefrontal network underpinned in part by the pre-commissural fornix.

## 1. Introduction

The fornix is the major white matter tract linking the hippocampal formation with distal brain sites (Saunders & Aggleton, 2007). Decades of neuropsychological evidence reveals that the integrity of the fornix is critical for episodic memory in humans and flexible spatial memory in rodents and primates (Gaffan, 1992; Aggleton & Brown, 1999; Aggleton et al., 2023).

While the fornix interconnects the hippocampal formation with numerous subcortical and cortical structures (Aggleton et al., 2010; Saunders & Aggleton, 2007), longstanding accounts of the neural substrates of episodic memory emphasise the importance of the direct hippocampal (HPC) projections to the anterior thalamic nuclei (ATN), along with the indirect fornical projections via the mammillary bodies (MB) to the ATN (Gaffan, 1992; Aggleton & Brown, 1999). According to Aggleton & Brown (1999) (see Aggleton et al., 2023 for subsequent developments) a common feature of anterograde amnesia is damage to any one of these structures, which collectively form an “extended hippocampal-diencephalic system” required for the encoding of new episodic information, binding the information to a spatial (and temporal) context, thus aiding subsequent recollection (Aggleton & Brown, 1999).

Direct tests of the functional unity of such a system have largely come from tests of allocentric spatial memory in rodents, informed by cognitive map theory (O’Keefe & Nadel, 1978), which posits that a fundamental function of the hippocampus (HPC) is the construction and maintenance of allocentric spatial maps of the environment, providing a spatial (and temporal) context that ‘scaffolds’ episodic memories (O’Keefe & Nadel, 1979; for an alternative account, see Eichenbaum et al., 1999). Studies using tasks that tap allocentric spatial processing, such as T-maze place alternation, or the Morris water maze (MWM) (Morris et al., 1982), have revealed that normal performance depends on the integrity of the HPC, fornix, MB, and ATN (Aggleton & Brown, 1999; Aggleton & O’Mara, 2022), consistent with the notion that the individual structures forming an extended hippocampal-diencephalic system function conjointly to support flexible spatial memory.

Likewise, HPC, fornix, MB, and ATN lesions result in equivalent levels of impairment on an object-in-place scene discrimination learning task in monkeys (Gaffan,1994; Parker & Gaffan, 1997a, 1997b; Murray et al., 1998). This task requires monkeys to learn and remember the location of a discrete object in a unique spatial scene and incorporates elements of both spatial and episodic-like memory (Gaffan, 1994). Equivalent levels of impairment on this task are seen in humans with fornix damage (Aggleton et al., 2000).

Complementing these findings, studies using diffusion-weighted magnetic resonance imaging (dMRI) in humans have found associations between inter-individual differences in fornix microstructure and inter-individual differences in scene recollection (Rudebeck et al., 2009), the spatio-temporal detail of autobiographical memories (Hodgetts et al., 2017), and scene discrimination learning (Postans et al., 2014).

While most lesion and dMRI studies have typically treated the fornix as a unitary tract, it is a complex fibre pathway with two main parts. The fornix bifurcates at the anterior commissure (AC) into a post-commissural (pocfx) and a pre-commissural (precfx) fornix segment (Aggleton et al., 2010). Most fornix fibres pass caudal to the AC to form the pocfx. These fibres originate in the subicular complex and terminate into the medial MB and en route also convey fibres into the ATN (Aggleton et al., 2010; Christiansen et al., 2016b). Other fibres, which originate in both the HPC and the subiculum, pass rostral to the AC as the precfx. These fibres travel anteriorly to terminate in the septal nuclei, nucleus accumbens (NAcc), and medial prefrontal cortex (mPFC) (Poletti & Creswell,1977; Jay & Witter, 1991; Cenquizca & Swanson, 2007).

While a few studies have attempted to compare the impact of selective pocfx versus precfx lesions on learning and memory (Henderson & Greene, 1977; Thomas, 1978; Tonkiss et al., 1990), it has proven challenging to confine damage to the target tract. An alternative approach is to use diffusion MRI tractography to isolate pocfx fibres (passing behind the AC) from precfx fibres (passing in front of the AC) (see Figure 2, method) (Christiansen et al., 2016a). Using this approach, in a sample of healthy older adults, Christiansen et al. (2016a) found a correlation between pocfx, but not precfx, microstructure and individual differences in visual recall from the Doors and People test (Baddeley et al., 1994). Similarly, Coad et al. (2020) found an association between pocfx, but not precfx microstructure and paired-associate object-in-location learning (PAL) (Lee et al., 2002). Together, these findings further support the importance of hippocampal-diencephalic connections in episodic and spatial memory.

Despite the emphasis on the direct fornical projections to the ATN, along with the indirect fornical projections to ATN via the MB, in current accounts of episodic and spatial memory, there are indications that other HPC connections conveyed via the fornix may play an important role in both episodic memory and spatial learning. In particular, regions of mPFC, directly connected (ipsilaterally) with the HPC (subiculum/CA1) by the precfx in both rodents (Jay & Witter, 1991; Cenquizca & Swanson, 2007) and primates (Barbas & Blatt, 1995; Aggleton et al., 2015), may be important for aspects of flexible goal-directed episodic and spatial memory in both rodents (Euston et al., 2012; Eichenbaum et al., 2017) and humans (Murray et al., 2017; Patai & Spiers, 2021).

One line of evidence comes from the study of allocentric spatial learning in tasks including the MWM. As with HPC lesions (Morris et al., 1982), fornix-transected rodents are impaired in learning this task when they are required to navigate flexibly from multiple start positions, but not when learning to navigate along a single route (Eichenbaum et al. 1990; Compton et al., 1997). An identical pattern of impairment is seen following lesions of mPFC (Compton et al., 1997; see also Sutherland et al., 1982). Disconnecting the two structures by silencing the HPC of one hemisphere and the mPFC of the contralateral hemisphere (‘crossed lesions’) mimics this impairment (Wang & Cai, 1998; see also Churchwell et al., 2010), indicating that goal-directed memory tasks seem to require intact functions of both hippocampus and mPFC and their functional interactions (for additional findings and reviews see Eichenbaum et al., 2017; Poucet & Hok, 2017; Ito, 2018).

Further evidence for the importance of HPC-mPFC interactions in spatial navigation in humans comes from fMRI studies, which have shown increased HPC-mPFC functional coupling during the adoption of an allocentric spatial virtual reality (VR) navigation strategy (Dahmani & Bohbot, 2015) and between subiculum and mPFC during VR navigational planning (Brown et al., 2016), as well as reduced HPC-mPFC functional coupling in individuals with developmental topographical disorientation (Iaria et al., 2014), a selective orientation deficit characterised by an inability to form cognitive maps. More recently. using lesion network mapping, Roseman et al. (2024) found that bilateral HPC and mPFC are key nodes in a network defined by resting-state functional connectivity (RSFC), damage to which impairs topographical navigation.

While these functional interactions likely reflect, at least in part, the influence of HPC on mPFC, neither crossed lesion disconnections nor fc-MRI can determine a direction of effect when two target sites are reciprocally linked. Both HPC and mPFC receive and send projections to multiple cortical and subcortical areas, and so there are many pathways by which the hippocampus and PFC could functionally interact, beyond the direct HPC-to-mPFC connections via the precfx. These routes include relays via parahippocampal cortices and via subcortical sites, including the nucleus reuniens and the anterior thalamic nuclei (Prasad & Chudasama, 2013; Anderson et al., 2016; Eichenbaum, 2017; Dolleman-van der Weel et al., 2019). It remains unclear then, whether the precfx plays an important role in flexible spatial learning.

In a prior publication (Hodgetts et al., 2020), we provided initial evidence for the involvement of the human fornix in spatial learning, showing that microstructure of the fornix (reconstructed as a single tract) in healthy young adults was related to inter-individual differences in learning to navigate to a hidden target location from multiple starting positions in a desktop-based VR analogue of the MWM task that is also sensitive to HPC lesions (Kolarik et al., 2018). Notably, the association between fornix microstructure and learning held when controlling for HPC volumes, which were not related to learning (for wider discussion see Weisberg & Ekstrom, 2021). Here, we present a novel analysis of this dataset, building on our prior publication in two ways. First, based on the protocol of Christiansen et al. (2016a) we used dMRI tractography to separately reconstruct the fibres forming the pocfx and those forming the precfx, and evaluated their microstructure based on the two-compartment free water elimination model, which enables more accurate estimation of diffusion properties of tissue by mitigating the partial volume effects caused by free water (Pasternak et al., 2009).

Second, advancing our previous analysis of virtual navigation learning, which used latency to find the hidden target location as a performance metric, we adapted a proximity-to-goal measure of navigational efficiency previously used in studies of the MWM (Gallagher et al., 2015; see methods for details). Proximity – and related spatial precision-based measures – are more sensitive to individual differences in MWM learning in rodents (Gallagher et al., 2015) and to the impact of HPC damage on desktop-based virtual navigation in humans (Kolarik et al., 2018). Again, consistent with studies of the MWM (Gallagher et al. 2015; Tomas Pereira & Burwell, 2015), we used a curve-fitting approach to compute a sensitive single index of spatial learning rate based on change in proximity-to-goal (i.e., reduction in search error) across learning trials. Using these more anatomically- and behaviourally-precise measures, we examined the relationship between microstructure of the pocfx and precfx and learning rate, providing novel insights into the fornical connections underpinning flexible and/or precise allocentric spatial learning in humans.

## 2. Method

### 2.1 Participants

Thirty-three healthy adult volunteers (15 men, 18 women; mean age = 24 years, range = 19-33 years) gave written informed consent to participate in this study. Portions of this data have been published previously (Hodgetts et al., 2020). Here we address a novel and distinct question, combining a new analytic approach to measure the precision of spatial learning, and a novel anatomically informed tractography protocol for reconstructing distinct fornix divisions. Participants were scanned at the Cardiff University Brain Research Imaging Centre (CUBRIC) and completed a VR analogue of the MWM in a separate cognitive testing session. All participants were fluent English speakers with normal or corrected-to-normal vision. The study was approved by the Cardiff University School of Psychology Research Ethics Committee.

### 2.2 Desktop VR navigation task

This article constitutes a novel analysis of data from a previously published study (Hodgetts et al. 2020), with detailed methods reported here in line with the recommendations of Thornberry et al. (2021).

Participants completed the virtual navigation task reported by Kolarik et al. (2018). The task was implemented using Unity 3D (Unity Technologies, San Francisco) and required participants to use the arrow keys on a keyboard to explore a 3D virtual art gallery from a first-person perspective (see Figure 1 for task schematic).

**Figure 1.**
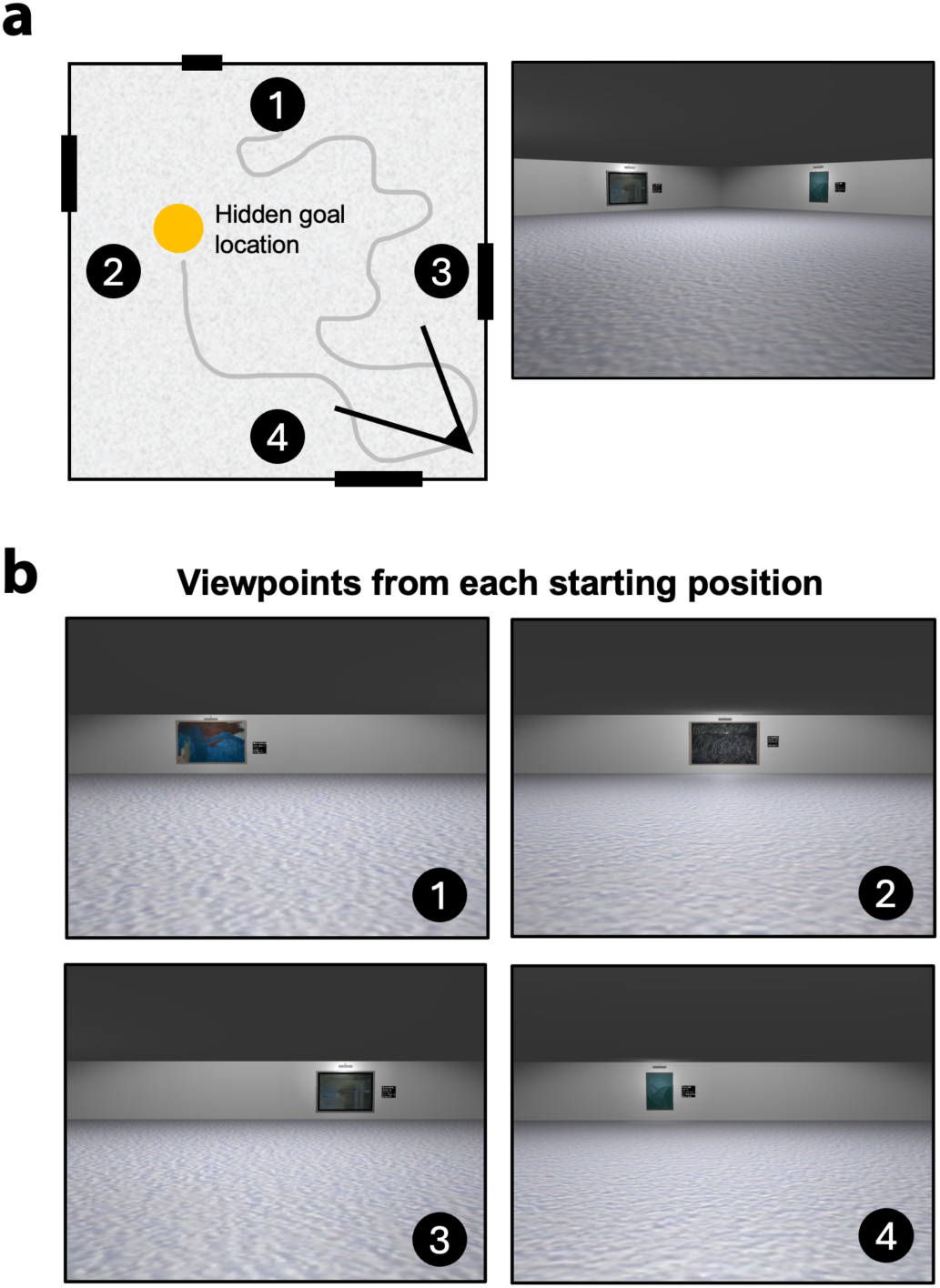
The desktop VR navigation task. (A) Bird’s-eye schematic and corresponding first-person view of the virtual art gallery arena that participants were required to navigate. Each wall of the arena features a painting that served as a ‘landmark’. Points 1, 2, 3, and 4 correspond to the arbitrary North, West, East, and South starting positions, respectively. (B) First-person viewpoints from each of the four starting positions, showing the painting landmarks used in the task.

The gallery room was 8 x 8 virtual metres, with 4 unique artworks, unevenly spaced, one on each wall, acting as landmarks. On a given trial, participants were required to locate a hidden sensor on the floor as quickly as possible. The hidden target was a .4 x.4 virtual metre square, occupying 0.25% of the total room area, whose position was fixed across consecutive trials. When participants’ search trajectory intersected the hidden sensor, it became visible and the caption “You found the hidden sensor” was shown in the centre of the screen.

Upon finding the sensor, a 10-second countdown appeared in the top left corner of the screen, which allowed participants to freely navigate and check their location with respect to the landmarks (the artworks) in the room before the next trial commenced. Following the countdown, an inter-trial screen appeared, and participants were able to click on a button to commence the next learning trial.

The maximum ‘free-search’ duration of each trial was 60 seconds. If a participant did not find the sensor within this timeframe, it became visible to allow the participant to move to and end the trial at the target location. Where this occurred, participants’ position and cumulative proximity to the target (see below) continued to be tracked during this additional ‘cued-navigation’ period.

The task involved 20 learning trials, which were divided into five blocks of four trials. In each of these blocks, participants started the trial from one of four possible starting positions (arbitrary North, South, East, West). Participants began from the North on trials 1, 7, 10, 16 and 19; the South on trials 4, 6, 9, 13 and 18; the West on trials 3, 8, 11, 14, and 17; and the East on trials 2, 5, 12, 15 and 20.

Each participant’s search trajectory was automatically recorded to a text file as Cartesian coordinates, with a frequency of 20 Hz.

### 2.3 MRI acquisition

Imaging data were acquired using a 3T GE Signa HDx MRI scanner (GE Healthcare) with an 8-channel receive-only head coil, at Cardiff University’s Brain Research Imaging Centre (CUBRIC). A standard T1-weighted 3D FSPGR sequence (178 axial slices, 1 mm isotropic resolution, TR/TE = 7.8/3.0s, FOV = 256 × 256 × 176 mm, 256 × 256 x 176 data matrix, 20° flip angle) provided high-resolution anatomical images.

A diffusion weighted single-shot spin-echo Echo-Planar Imaging (EPI) pulse sequence was used to acquire whole-brain dMRI data (60 contiguous slices acquired along an oblique-axial plane with 2.4 mm thickness and no gap, TE = 87 ms; voxel dimensions = 2.4 × 2.4 × 2.4 mm^3^; FOV = 23 × 23 cm^2^; 96 × 96 acquisition matrix). The acquisition was cardiac gated, with 30 isotropic directions at b = 1200 s/mm^2^. In addition, three non-diffusion weighted images were acquired with b = 0 s/mm^2^.

### 2.4 Diffusion MRI preprocessing

DWI preprocessing, including correction for participant head motion and Eddy currents, was performed with ExploreDTI v 4.8.3 (Leemans et al., 2009). We used custom MATLAB scripts to perform free water elimination (FWE) based on a bi-tensor model to separate isotropic (free water) and anisotropic (tissue) diffusion compartments (Pasternak et al., 2009). This FWE pipeline includes spatial regularization (since estimation of the model is ill-posed in single shell dMRI) (Pasternak et al., 2009). FWE mitigates the adverse impact of CSF partial volume on diffusion tensor (DTI) metrics. This is especially crucial when examining the fornix, which borders the 3rd ventricle (De Santis et al., 2014).

Following FWE, corrected voxel-wise DTI indices were computed. Fractional Anisotropy (FA) – a dimensionless DTI-based index proposed to reflect axonal organization (Basser, 1997), represents the extent to which diffusion within biological tissue is anisotropic (restricted along a single axis) and can range from 0 (isotopic diffusion) to 1 (anisotropic diffusion). The resulting FA maps were inputs for tractography.

### 2.5 Tractography

Deterministic tractography was performed from all voxels based on constrained spherical deconvolution (CSD) (Jeurissen et al., 2011). CSD allows for the representation of bending/crossing/kissing fibres in individual voxels, as multiple peaks in the fibre orientation density function (fODF) can be extracted within each voxel (Dell’Acqua and Tournier, 2019). The step size was 1 mm, and the fODF amplitude threshold was 0.1. An angle threshold of 30° was used to prevent the reconstruction of anatomically implausible fibres.

To generate 3D fibre reconstructions of each tract segment, waypoint region-of-interest (ROI) gates were drawn manually onto whole-brain FWE-corrected FA maps. The waypoint ROIs defined the tracts based on a ‘SEED’ point and Boolean logical operations: ‘NOT’ and ‘AND’. The ‘NOT’ and ‘AND’ gates corresponded to whether tracts passing through were omitted from analyses or retained, respectively. These gates were combined to reconstruct the tracts, based on anatomical plausibility (Schilling et al., 2020). Initially, a multiple ROI approach was applied to reconstruct the fornix and subsequently fornix tract division was performed.

#### 2.5.1 Fornix reconstruction

The fornix was initially reconstructed in its entirety using a multiple-ROI approach (Hodgetts et al., 2017). A ‘SEED’ ROI encapsulating the fornix body was drawn on a coronal section. This was combined with an ‘AND’ ROI drawn on an axial section immediately inferior to the splenium of the corpus callosum, with an area sufficient to capture the crus fornici.

Exclusionary ‘NOT’ ROIs were drawn on an axial section covering the corpus callosum body, and on coronal sections covering the whole brain immediately anterior to the fornix pillars and posterior to the crus fornici. Post hoc exclusionary ‘NOT’ ROIs were drawn to exclude any extant spurious streamlines as required. The anterior body of the fornix was then split into its pocfx and precfx column segments (Figure 2).

**Figure 2.**
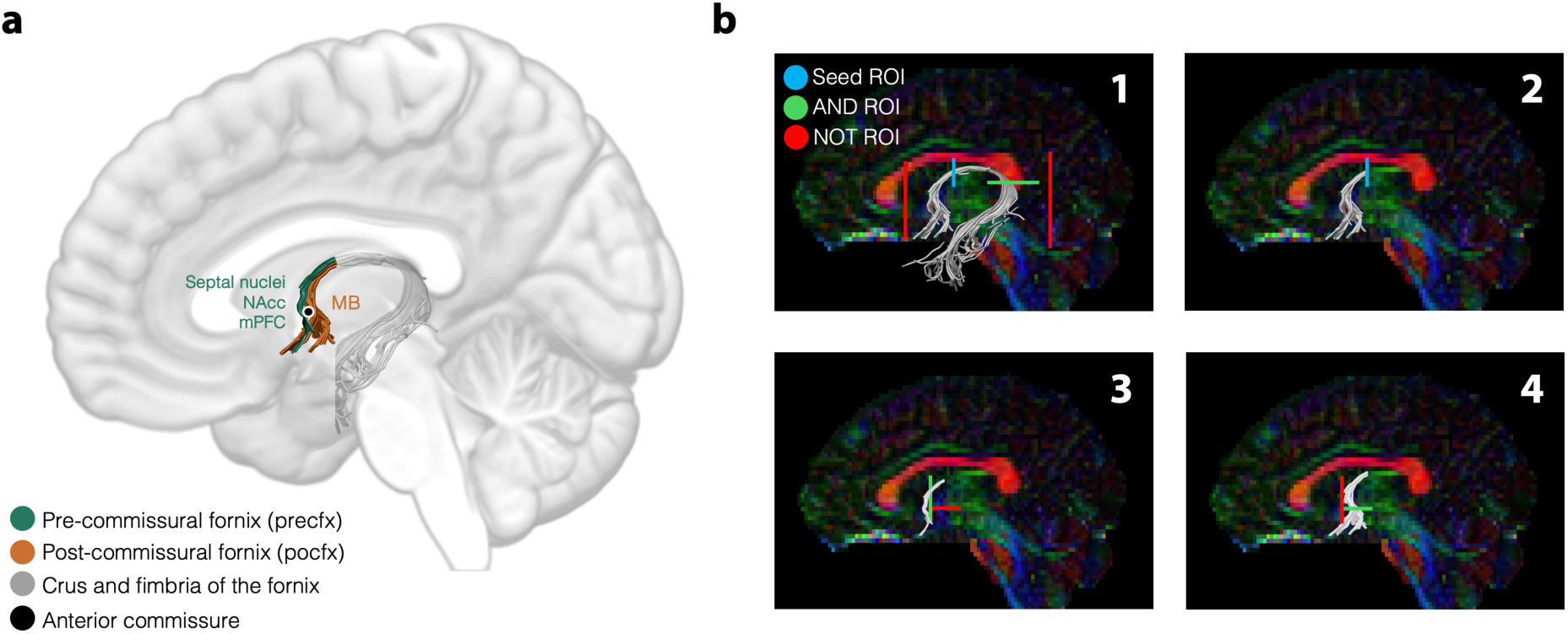
Pre- and post-commissural fornix segmentation. (A) Schematic illustration of the anatomical landmarks used to differentiate pre- and post-commissural fornix segments (shown in green and orange, respectively). The key termination sites of the precfx and pocfx based on anatomical tract tracer studies are shown beside each tract segment in their respective colour (see main text; mPFC = medial prefrontal cortex; MB = mammillary bodies; NAcc = nucleus accumbens). (B) Approach and waypoint ROIs used for reconstructing the precfx and pocfx in ExploreDTI (see Sections 2.5.1 and 2.5.2 for further detail).

#### 2.5.2 Pre- and post-commissural fornix segmentation

The anterior body of the fornix was first extracted from the fornix reconstructions using the ‘splitter’ tract segmentation tool within ExploreDTI (see Figure 2), by drawing an ‘AND’ ROI immediately anterior to the existing ‘SEED’ ROI (see previous section), at the point where the tract descends towards the crus fornici. This step removed the crus and fimbria components from the fornix reconstructions, and was performed to minimise the risk of partial volume effects in the final segmentations due to an intermingling of pocfx and precfx streamlines as they progress into and beyond the crus fornici (Saunders & Aggleton 2007).

The anterior body of the fornix was then further divided into its precfx and pocfx segments. The precfx was extracted by drawing an additional ‘AND’ ROI on a coronal section at the level of the AC, and an exclusionary ‘NOT’ ROI that intersected this ‘AND’ ROI on an axial section. To reconstruct the pocfx, the placement of these ROIs was swapped. Fibre streamlines were therefore only included in the precfx segmentations if they extended anterior to the AC, whereas the pocfx segmentations retained only those streamlines that descended posterior to the AC (see Figure 2 for representative reconstructions).

### 2.6 Search trajectory and spatial learning analyses

In our earlier publication (Hodgetts et al., 2020), performance on each learning trial was defined by the time (in seconds) to reach the hidden target sensor. Latencies to find a target location are, however, a relative imprecise measure of place learning (Gallagher et al., 2015).

To increase sensitivity to inter-individual differences in spatial learning, for each learning trial, participants’ proximity to the centre of the hidden sensor was sampled at 1-second intervals for the duration of their search, and this data was summed to compute a *cumulative proximity* measure (Gallagher et al., 2015). Participants whose search area is concentrated near the sensor will have a lower cumulative proximity score (i.e., smaller cumulative search error) compared to individuals that spend an equivalent time searching for the sensor but within a search area that is more widely distributed around the virtual room (see Supplementary Figure S1). A lower score indicates that the subject’s path is more direct and efficient toward the goal.

For each trial, participants’ initial proximity to the sensor was subtracted from this cumulative proximity measure, since the 4 possible starting locations were non-equidistant from the target sensor. Prior research has demonstrated that proximity measures are more sensitive indices of allocentric place learning compared to latency metrics (Gallagher et al., 2015; Tomas Pereira & Burwell, 2015; Zhong et al., 2017).

We then used a curve-fitting technique to derive a single participant-specific index of learning rate from these trial-wise cumulative proximity measures (for a similar approach in rodent studies of the MWM see Gallagher et al., 2015; Tomas Pereira & Burwell, 2015). Here, a learning curve was fit to subject-specific trial-wise cumulative proximity measures using a power function:

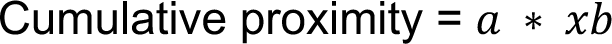

The slope parameter (*b*) indicates the slope curvilinearity in individual subjects, with lower, negative values, reflecting a more rapid descent from start to minimum search error (i.e., a faster learning rate).

As discussed in Hodgetts et al. (2020; see Commins et al., 2023 for an alternative approach to addressing this issue), simple curve-fitting techniques can, however, result in counterintuitive results when applied across all learning trials, such that some of the fastest learners will present the poorest model fits. For instance, this model provides a poor fit for individuals who learn the task quickly and plateau, but do not then sustain this level of performance until the learning trial series is completed (see example data from two participants in Figure 3A; see Supplementary Figure S2 for all learning curves). This pattern of performance may reflect cognitive/motivational factors (e.g., mind-wandering, novelty seeking) affecting performance in trials towards the end of the task. To account for this complexity in learning patterns, we therefore used a data-driven approach to identify and apply a cut-off point to each individual participant’s trial-wise performance data. A second-order polynomial model was fitted to everyone’s trial-wise performance data using the Matlab curve-fitting toolbox (Mathworks, Inc.). The participant-specific cut-off was defined as the ‘trough’ of the resulting curve, at which point the first derivative of the second-degree polynomial crosses zero (Figure 3, middle). All trials up to and including this participant-specific trial cut-off point were modelled using the power function described above (modal number of trials included = 13; range = 10 - 16). Critically, as shown in Figure 3B, faster learning rates were observed for the truncated model compared to the model derived from all trials, and this difference was statistically significant (t(32) = −7.39, P < 0.001, d = −1.07). Further, the group location/trajectory heatmap for the participant-specific ‘cut-off’ trial illustrates that participants’ searches converged around the target location and were not as randomly distributed around the arena compared to trial 1, consistent with successful place learning by the ‘cut-off’ trial (Figure 3C).

**Figure 3.**
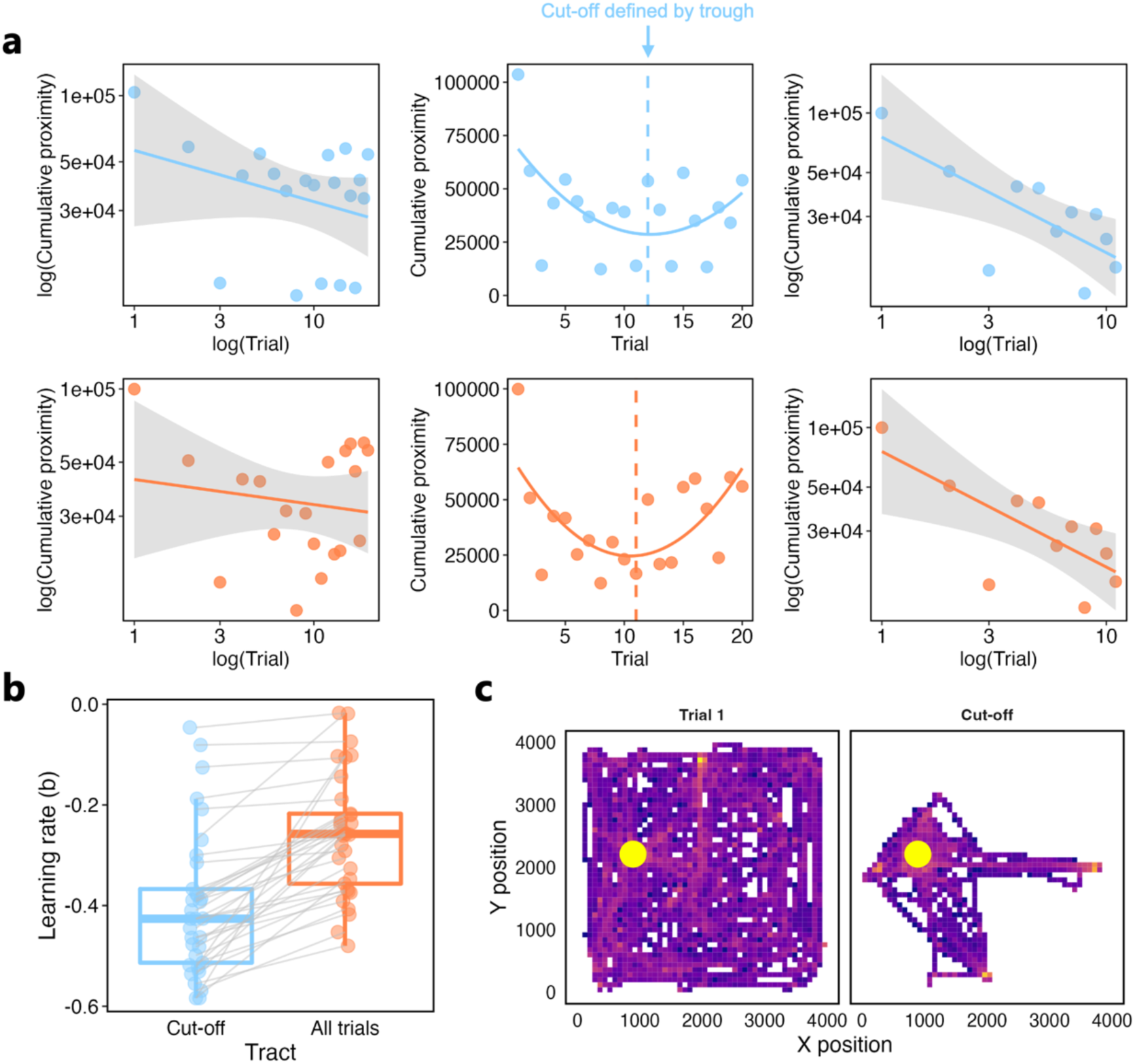
The method used to determine the number of trials to-be-modelled in each participant. (A) Navigational learning data from two example participants are shown. Both display early learning followed by performance plateau and then poorer / more variable performance in later trials. A power function fits this pattern of cumulative proximity data poorly (left). To address this, a second-order polynomial was fitted to the trial-wise cumulative proximity data for each participant. The trial at which the first derivative of this polynomial crossed zero (i.e., the trough of the fitted curve) was taken as the point of minimum search error, and thus used to define the number of trials to be modelled (middle). Trials up to this participant-specific cut-off were then fit with a power function, and the *b* parameter was derived to index learning rate (right). Power fits were obtained by linearly fitting the log-transformed data. (B) Boxplot comparing values of the *b* parameter derived when using all trials (as in A, left) versus trials up to the cut-off (as in A, right). A statistical comparison of these fits can be found in the main text. (C) Group-level trajectory heatmaps illustrating participants’ search paths relative to the hidden target (yellow circle) on Trial 1 and at the cut-off trial. On Trial 1, trajectories were widely dispersed across the arena; by the cut-off trial, search behaviour was more focused around the hidden sensor. Trajectories are collapsed across participants for visualisation purposes.

### 2.7 Statistical analysis of fornix microstructure-learning rate associations

Statistical analyses were conducted in R (R Core Team, 2021).

Since higher FA is considered indicative of increased myelination, and improved organisation, cohesion and compactness of white matter fibre tracts (Beaulieu 2002), and since lower *b* values indicate faster learning (a more rapid decrease in cumulative search error across trials), we predicted a negative association between fornix FA and the learning rate parameter (*b*). We therefore computed directional Pearson correlations between FA of the precfx and pocfx and learning rate, *b* (Greenland et al., 2016). As we computed correlations across two tract segments, we applied a Bonferroni correction to our significance threshold, so that correlations with p ≤ 0.025 were considered statistically significant.

Additional Bayesian correlation analyses were conducted using JASP (Version 0.19.3) (JASP Team, 2024). From this, we report default Bayes factors and 95% Bayesian credible intervals. The Bayes factor (BF) grades the intensity of the evidence that the data provide for the alternative hypothesis (H1) versus the null (H0) (or vice versa) on a continuous scale (Jeffreys, 1961; Wagenmakers et al., 2016).

Pearson correlations between learning rate and precfx and pocfx FA respectively were compared using a directional Steiger Z-test, computed using R package ‘cocor’, (Diedenhofen & Musch 2015).

Prior to these analyses, we identified and excluded participants whose cross-trial performance data did not provide robust evidence that place learning had occurred, for example, those not engaging with the task or those participants who randomly located the target very quickly on the first trial and who therefore had little opportunity to further improve performance across trials (i.e., ‘non-learners’). This was achieved using a resampling approach in which an individual participant’s cross-trial performance data was shuffled over 500 permutations to derive confidence intervals (CIs). Participants with a model R^2^ falling outside the 68% CI (i.e., 1 SD) of their participant-specific random distribution were marked for exclusion from correlational analyses. Nine individuals were thus excluded, and a two-tailed Welch’s unequal variances t-test confirmed that the learning rates for these ‘non-learners’ (mean = −0.223, SD = 0.127) were significantly poorer than those derived from the retained participants (mean = −0.469, SD = 0.075; t (32) = −6.919, p < 0.001, d = −2.36). All subsequent analyses (including comparisons between tract segments) focused on those ‘learner’ participants whose trial-wise performance data provided robust evidence of successful place learning.

### 2.8 Data and code sharing

Anonymised output data and the code used within this project are freely available at https://osf.io/vsk3u/ ensuring that the figures and values reported in this manuscript are reproducible. However, ethical restrictions, relating to General Data Protection Regulation, do not allow for the public archiving of the raw study data. Access to pseudo-anonymised data could be granted after signing and approval of data transfer agreements. For this, readers should contact Dr Carl Hodgetts and Prof. Andrew Lawrence.

## 3. Results

### 3.1 Comparing fornix segment microstructure

We derived measures of participants’ learning rates (*b* values; mean = −0.47, SD = 0.075) as well as FWE-corrected tract-averaged FA of the precfx (mean = 0.40, SD = 0.038) and pocfx (mean = 0.365, SD = 0.0446). FA of the precfx was significantly greater than that of the pocfx (t(23) = 3.223 p < 0.004, d = 0.831) (replicating the findings of Coad et al., 2020).

A two-tailed Pearson correlation revealed that precfx and pocfx FA measures were not significantly correlated (r(22) = 0.206, p = 0.335, BF_01_ = 2.545, 95% [-0.204, 0.536] (Figure 4).

**Figure 4.**
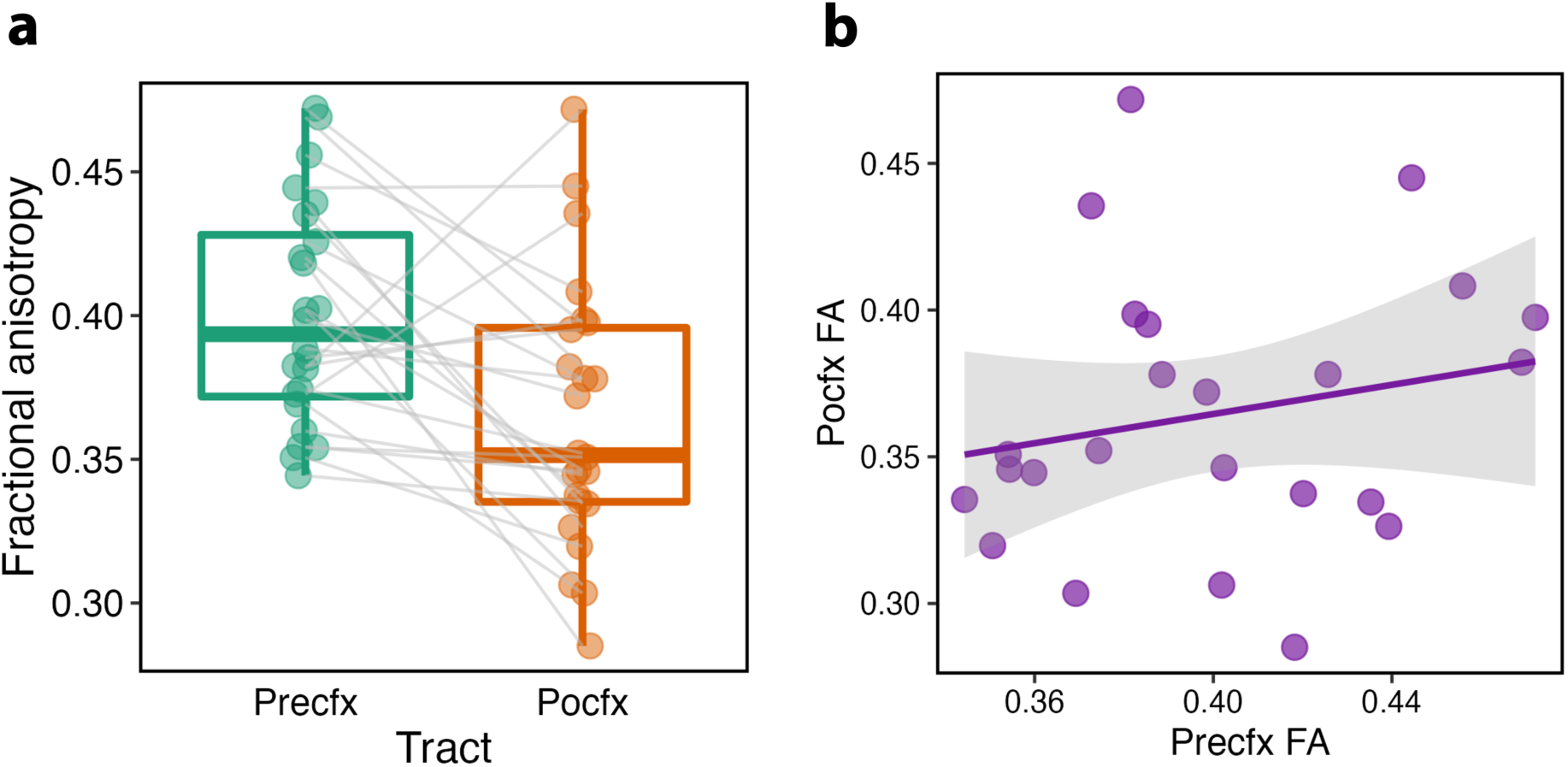
Comparing fornix segment microstructure. (A) Boxplot comparing fractional anisotropy (FA) values for the precfx and pocfx, with individual data points shown. (B) Association between precfx and pocfx FA. The grey shaded area represents the 95% confidence interval.

Importantly, there was similar inter-individual variability in FA values across the two fornix segments (Pitman-Morgan test for (non)equality of variances (t (31) = 0.570, p = 0.573) (Figure 4; computed using the ‘PairedData’ R package, Champerly, 2018).

### 3.2 Correlation between fornix segment microstructure and spatial learning

For our planned analyses, we ran directional Pearson correlations between participants’ learning rate (*b*) and fornix microstructure. These revealed a significant negative association between participant-specific learning rates and precfx FA (r(22) = −0.443, p = 0.015, BF_-0_ = 4.591, 95% CI [-0.079, −0.696], such that participants with higher precfx FA had faster learning rates. By contrast, learning rate was not significantly correlated with pocfx FA (r(22) = 0.037, p = 0.569, BF_0-_ = 4.485, 95% CI [-0.006, 0.411]). Furthermore, the correlation between learning rate and precfx FA was significantly greater than that with pocfx FA (z = −1.884, p = 0.033; Figure 5).

**Figure 5.**
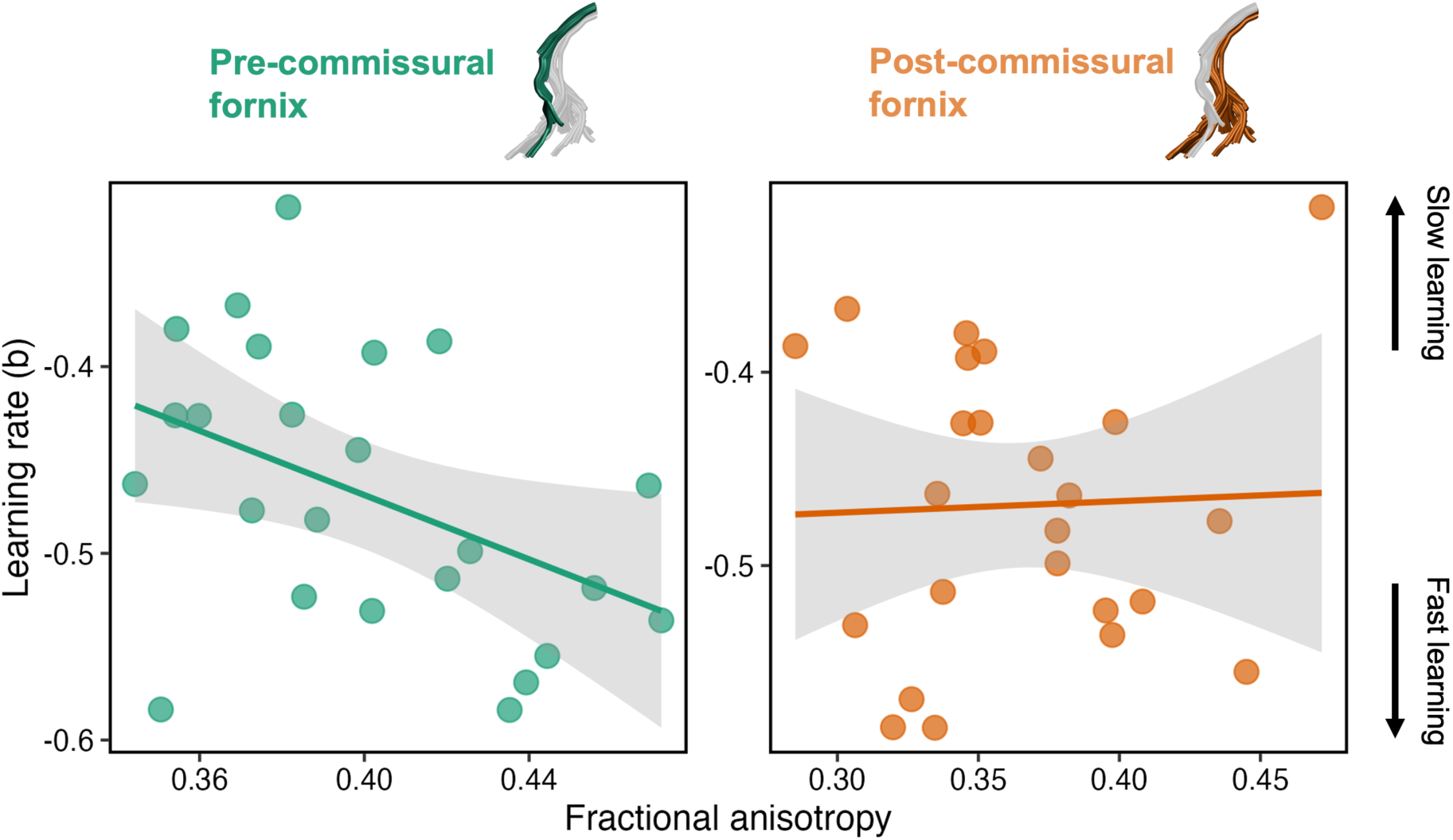
The association between navigational learning rate (*b*) and precfx (left, green) versus descending pocfx fornix FA (right, orange). The grey shaded area in each plot represents the 95% CIs.

### 3.2 Post hoc analysis: influence of sex on microstructure-learning correlations

Based on prior findings of sex differences in spatial navigation (including desktop VR MWM performance), typically characterised by a performance advantage for men, on average (Astur et al., 1998; Moffat et al., 1998; Woolley et al., 2010), we conducted a post hoc analysis to examine whether our main findings remained when controlling for sex (9 men, 15 women). The significant negative correlation between precfx FA and learning rate held when controlling for sex (r = −0.471, p = 0.024). Further, we observed no significant difference between men (mean = −0.494, SD = 0.081) and women (mean = −0.454, SD = 0.069) on our learning rate parameter (t(22) = 1.227, p < 0.215, d = 0.538).

## 4. Discussion

The fornix carries numerous projections both to, and from, the hippocampal formation, raising the fundamental question of which fornical pathways are most critical for spatial learning and memory (Aggleton et al., 2010). Here, we applied an anatomically guided diffusion tractography protocol to separately reconstruct pre-commissural (precfx) and post-commissural (pocfx) fornix fibres in vivo (Christiansen et al., 2016a) and evaluate their microstructure, indexed by free water corrected fractional anisotropy (FA), in a healthy young adult sample. To capture inter-individual differences in spatial learning performance, we used a continuous, graded measure based on cross-trial improvement in proximity-to-target location in a desktop VR analogue of the MWM (Kolarik et al., 2018). This measure considers distance-to-the-goal along search trajectories and is more sensitive to relational, allocentric spatial strategy relative to search latencies alone (Gallagher et al., 2015). Using this approach, we found that inter-individual variation in precfx, but not pocfx, microstructure was significantly correlated with spatial learning, such that participants with higher FA in the precfx had faster learning rates, reflecting steeper cross-trial reductions in cumulative search error during learning. Our findings highlight, for the first time, the importance of one specific set of HPC connections for efficient flexible spatial learning in humans.

Anatomical tract tracing studies in rodents and primates show that the precfx principally innervates the septal nuclei (including medial and lateral septum), NAcc, and mPFC (Swanson & Cowan, 1977; Jay & Witter, 1991; Saunders & Aggleton, 2007; Aggleton et al., 2015). The precfx also provides a modest source of cholinergic input to the HPC from the medial septum-diagonal band complex (MSDB) (MacLean et al., 1997). Precfx fibres do not, however, appear to innervate the diencephalon (Mathiasen et al., 2019). Since lesions of septal nuclei, NAcc and MSDB rarely result in impairments in allocentric spatial learning (Rawlins & Olton, 1982; Baxter et al., 1995; Floresco et al., 1996), our findings likely highlight the importance of connectivity between the HPC and mPFC mediated by the precfx in efficient and/or precise allocentric spatial learning.

While HPC place cells have long been established as the basis for a detailed representation of current location relative to the overall spatial layout, forming the elements of a cognitive map of space (O’Keefe & Nadel, 1978; O’Keefe & Krupic, 2021; Ekstrom et al., 2003; Hori et al., 2005; but see Eichenbaum et al., 1999 who argue that ‘place’ cells represent the set of experiences within a particular location), increasing evidence from rodent and human lesion and electrophysiological studies highlights a central role for the mPFC in goal representation, route planning, and top-down control of navigation (Ekstrom et al., 2003; Euston et al., 2012; Poucet & Hok, 2017; Patai & Spiers, 2021). Furthermore, both fMRI studies in humans (reviewed in Patai & Spiers, 2021) and disconnection studies in rodents (reviewed in Eichenbaum, 2017) show that functional interactions between HPC and mPFC are critical for optimal performance on certain goal-directed memory tasks. In the domain of goal-directed spatial navigation, several recent models incorporate both HPC and mPFC into complex bidirectional circuitry in which map-based route planning requires finely coordinated interactions between ‘bottom-up’ spatial/contextual representations generated in the HPC and ‘top-down’ task/goal representations in the mPFC (Shapiro et al., 2014; Ito, 2018, Dolleman-van der Weel et al., 2019).

There are several pathways that could support HPC-mPFC functional interactions. One pathway for interaction involves a direct projection from the HPC to the mPFC, and there are other indirect pathways through intermediary thalamic and rhinal cortical routes that are bidirectional (Prasad & Chudasama, 2013; Anderson et al., 2016; Aggleton & O’Mara et al., 2022; Messanvi et al., 2023). Critically in relation to our findings, the direct anatomical connectivity from HPC to mPFC is conveyed by the precfx. In rodents, the entire longitudinal extent of the subiculum/CA1 is connected – via the precfx – with the mPFC, with connectivity increasing progressively in strength from dorsal to ventral hippocampus (Cenquizca & Swanson, 2007; Jay & Witter, 1991; Hoover & Vertes, 2007; Ährlund-Richter et al., 2019).

Similarly in primates, the precfx provides the exclusive route for subiculum/CA1 projections to mPFC (Aggleton et al., 2015; Barbas & Blatt, 1995; Carmichael & Price, 1995), with relatively more projections arising from anterior HPC.

Recent findings challenge the traditional spatial-emotional dichotomy along the dorsal (posterior)-ventral (anterior) axis of the hippocampus (Moser & Moser, 1998), instead supporting a functional gradient, in line with the anatomical gradient described above. Lesion studies in rodents confirm that flexible spatial learning requires both the dorsal (dHPC) and ventral (vHPC) hippocampus (Avigan et al., 2020; Hwang et al., 2024) (for similar findings in humans, see Kolarik et al., 2018). It has been suggested that vHPC plays a unique role in in interfacing between dHPC place-based and mPFC goal-based representations (McKenzie et al., 2016). Consistent with this, lesions of vHPC (Blanquat et al., 2013; Babl & Sigurdsson, 2025) and optogenetic silencing of monosynaptic vHPC-mPFC connections (Spellman et al., 2015) impair mPFC goal-location and goal-value encoding, while dHPC lesions and silencing do not.

In rodents, oscillatory synchrony is a key feature of bidirectional functional coupling between HPC and mPFC (Hyman et al., 2011; Eichenbaum, 2017; see also Chrastil et al. 2022 for related findings in humans using scalp-recorded EEG during VR maze-learning). The direction of the HPC-mPFC projection via the precfx aligns with the HPC-leading-mPFC directionality of information flow during spatial encoding-related increases in theta and gamma coherence between these regions seen during neural recordings in rodents (Siapas et al., 2005; Spellman et al., 2015; Nardin et al., 2023), which appears to depend upon the integrity of the ventral hippocampus (O’Neill et al., 2013). Since variation in the microstructural properties of white matter tracts can influence timing and synchronization of oscillatory activity between distal brain regions (Pajevic et al., 2014; Bells et al., 2017, 2019), inter-individual differences in precfx microstructure may, therefore, impact the efficiency/fidelity of spatial information transfer from HPC to mPFC and hence the efficiency and/or precision of spatial learning.

Importantly, the precfx is not the only route by which spatial information can be transferred from HPC to mPFC. The nucleus reuniens (NRe) of the thalamus serves as a critical bidirectional relay between HPC and mPFC (Dolleman-van der Weel et al., 2019; Mathiasen et al., 2020; Messanvi et al., 2023) and has been argued to modulate HPC-mPFC theta coordination (Eichenbaum et al., 2017; Dolleman-van der Weel et al., 2019; but see de Mooij-van Malsen et al., 2023). Our findings are important therefore in clearly indicating that the direct HPC inputs to the mPFC via the precfx are key to the functioning of broader bidirectional HPC-mPFC circuitry underpinning spatial learning and memory.

Notably, direct anatomical HPC-mPFC connectivity via the precfx is one-way. There are no direct return projections from mPFC to HPC (Andrianova et al., 2023). Models of mPFC regulatory action upon the HPC emphasise the importance of indirect routes via the nRe, as well as rhinal cortices (Anderson et al., 2016; Eichenbaum, 2017; Dolleman-van der Weel et al., 2019). This circuitry is thought to enable executive control of HPC spatial representations to meet specific task demands. Experimental lesions and inactivation studies in rodents confirm the importance of this feedback: disrupting mPFC or nRe function impairs task-dependent HPC activity and flexible navigation (Shapiro et al., 2014; Ito et al., 2015; Dolleman-van der Weel et al., 2019). Collectively, this bidirectional circuit, in which the precfx is a key link, helps to integrate the HPC cognitive map and a prefrontal abstract task ‘map’ or schema (Van de Maele et al., 2024). Spatial Information flows, at least in part via the precfx, ‘bottom-up’ from HPC during initial encoding of location and context, then ‘top-down’ from mPFC to bias future navigational trajectories based on goals and learned task rules. Coordination between these ‘maps’ is critical for prospective planning, decision-making, and optimising cognitive flexibility (Shapiro et al., 2014; Poucet & Hok, 2017; Ito, 2018; Van de Maele et al., 2024).

Considering the emphasis placed on HPC-diencephalic connections mediated by the pocfx, but not the precfx, in longstanding accounts of episodic and spatial memory (Gaffan, 1992; Aggleton et al., 1999; Aggleton et al., 2023) it was somewhat surprising that we found evidence *against* an association between pocfx microstructure and learning rate. One important caveat is that our pocfx tract reconstructions principally involve the connections of the HPC with the MB and largely exclude the projections to the ATN, since these turn immediately caudally to project into the ATN as the fornix columns descend to reach the MB (Poletti & Creswell, 1977; Christiansen et al., 2016b). The direct ATN projecting fibres do not form a discrete tract, but appear to remain diffuse (Mathiasen et al., 2019) and difficult to reconstruct with dMRI tractography (Ozdemir et al., 2024). While lesion and crossed disconnection studies in rodents reveal the importance of these direct subicular-ATN connections for flexible spatial learning (Aggleton & O’Mara, 2022), including in the MWM (Warbuton et al., 2000), it is the case that complete fornix section can lead to greater impairments on certain tasks of spatial learning/memory (e.g., T-maze continuous alternation) than either ATN or MB lesions (Aggleton et al., 1995), further pointing to the importance of fornix connections beyond those linking the HPC with the diencephalon.

It might also be argued that a key difference between our findings and those in rodents and primates regarding the contribution of HPC-MB connections carried by the pocfx is that stationary desktop VR navigation does not provide idiothetic, self-motion cues (see Starrett & Ekstrom, 2018 for discussion). Gaffan (1998) proposed that idiothetic cues generated by self-movement – whether of the whole body, limbs, or eyes – could, in theory, participate in object-in-place memory, and that the HPC and fornix provide such an idiothetic signal. However, lesion studies in rodents fail to support the idea that the fornix is important in conveying idiothetic information (Bussey et al., 2000).

Strikingly, and consistent with our findings, Vann and colleagues (Vann, 2013; Vann et al., 2011) have reported that selective lesions to the descending pocfx in rats, which completely disconnect the subicular projections to the MB, but spare those with the ATN, as well as sparing connections mediated by the precfx, have little-to-no impact on spatial memory tests, including the MWM and T-maze place alternation, that are sensitive to MB, mammillothalamic tract, ATN, and HPC lesions. A clear implication of these findings (with the caveat that they represent a single dissociation) is that the direct HPC-MB connectivity mediated by the pocfx may be less critical, and the direct HPC-mPFC connectivity mediated by the precfx more critical, to flexible spatial learning than previously highlighted in accounts of an “extended hippocampal-diencephalic system” (Gaffan, 1992; Aggleton & Brown, 1999). Notably, we (Williams et al., 2020) also found a correlation between inter-individual differences in episodic autobiographical past and future thinking and precfx, but not pocfx, microstructure. Together, these findings highlight the potential importance of the direct HPC connections to mPFC carried by the precfx in the ability to flexibly form and utilise spatial/episodic representations in line with task demands (Eichenbaum, 2017; Murray et al., 2017).

Although our findings highlight the contributions of HPC connections conveyed by the precfx, this exercise is not exclusive, that is, demonstrating the importance of one set of connections does not negate the importance of other connections (Aggleton & O’Mara, 2022). Previous findings in humans link pocfx microstructure to other aspects of episodic memory (Christiansen et al., 2016a, Coad et al., 2020), including tests sensitive to MB pathology (Tsivilis et al., 2008), and there is clear evidence in rodents that subicular neurons targeting MB are critical for some forms of spatial memory (Li et al., 2025). Understanding the complementary roles of the hippocampal-diencephalic and hippocampal-basal forebrain/prefrontal networks mediated by the fornix in learning and memory is an important future goal (Aggleton & O’Mara, 2022; McNaughton & Vann, 2022).

The present study has limitations that should be born in mind. Although FA is highly sensitive to the microstructure of fibers (De Santis et al., 2014), it lacks biological specificity, and may reflect myelination, axonal diameter, axonal density, and fibre dispersion (Beaulieu, 2002). Future multi-modal investigations using multi-shell diffusion MRI and advanced biophysical modelling to estimate specific tract microstructural properties, alongside measures of functional (oscillatory) connectivity using electrophysiological imaging with MEG (Read et al., 2025) may provide further insight into the specific biological attributes and network interactions underlying the current microstructure-learning associations.

Furthermore, virtual tract renditions are created from estimations of water diffusion directionality, not from the anatomy itself, and characterization of fibres is limited by the MRI technology (e.g., higher gradient strengths enable dMRI acquisitions with higher spatial resolution, which enables the resolution of smaller white-matter bundles, Jones et al., 2018). Although we constructed virtual tract renditions using anatomical knowledge, it is not possible to test the specificity of the tractography methods, for each participant, without knowing the true underlying anatomy (Schilling et al., 2020).

While our sample size was comparable to related investigations (e.g., Brown et al., 2014; Chrastil et al., 2017), replicable and precise results are more likely when statistical power is high (Button et al., 2013). Critically, however, it is quite possible for low-power experiments to have high evidential value, and for high-power experiments to have low evidential value (Wagenmakers et al., 2015). Here, default BFs > 3 show that our findings provide substantial evidence, according to Jeffreys’s (1961) conventions, for a positive correlation between precfx microstructure and learning, and for a null correlation between pocfx microstructure and learning. That said, it will be important to extend our findings to larger samples, preferably using longitudinal and/or training designs that can examine the relationship between experience, brain microstructure, and spatial learning abilities (Coutrot et al., 2022).

In line with our hypothesis-driven analysis approach we utilised a spatial precision-based metric to assess learning. Further supporting the importance of this approach, a recent study found that individuals with fornix lesions showed intact schematic memory but impaired fine-grained detailed memory on sketch maps of recently and remotely experienced neighbourhoods and houses (Li et al., 2024). Nevertheless, we note that although our virtual maze task (VWM) and scoring protocol have strong validity (Gallagher et al., 2015; Kolarik et al., 2018), there are a plethora of VWM tasks and scoring protocols that are not standardised across research groups (see Thornberry et al., 2021 for discussion), making replication and generalization of experimental findings difficult. Future work in this area will benefit from adoption of better reporting standards, pre-registration of (standardised) protocols and scoring, and reproducible analysis pipelines.

In summary, we report a novel association between white matter microstructure of the precfx, but not pocfx, and inter-individual differences in flexible spatial learning. Together with similar findings in episodic memory (Williams et al., 2020), these findings help inform anatomical models of spatial and episodic memory and elucidate a potential anatomical substrate by which direct HPC-to-mPFC connectivity, as part of wider bidirectional HPC-mPFC circuitry, enable flexible encoding, retrieval, and recombination of spatial/episodic information in line with task demands (Eichenbaum, 2017; Murray et al., 2017).

## Additional information

### CRediT authorship contribution statement

Carl J. Hodgetts: conceptualization, formal analysis, funding acquisition, data curation, visualization, validation, writing – reviewing and editing. Mark Postans: conceptualization, formal analysis, methodology, writing – original draft, writing – reviewing and editing. Angharad N. Williams: formal analysis, writing – reviewing and editing. Kim S. Graham: supervision, funding acquisition, project administration, writing – reviewing and editing. Andrew D. Lawrence: conceptualization, formal analysis, funding acquisition, supervision, writing – original draft.

### Data availability

Anonymised output data and the code used within this project are freely available at https://osf.io/vsk3u/ so that the figures and values reported in this manuscript are reproducible. However, ethical restrictions, relating to General Data Protection Regulation, do not allow for the public archiving of the raw study data. Access to pseudo-anonymised data could be granted after signing and approval of data transfer agreements. For this, readers should contact Dr Carl Hodgetts and Prof. Andrew Lawrence.

## Acknowledgements

This work was supported by the Biotechnology and Biological Sciences Research Council (BBSRC) [BB/V010549/1 (CJH); BB/V008242/1 and BB/V008242/2 (ADL)]. We would like to thank Professor Arne Ekstrom (University of Arizona) for sharing the VR navigation task, Martina Stefani for assisting with data collection, Dr C. John Evans and Peter Hobden for scanning support, and Dr Greg Parker for providing the free water correction pipeline. For the purpose of open access, the authors have applied a creative commons attribution (CC-BY) license to any author accepted manuscript version arising.

