## Supplemental information for "Fornix subdivisions and spatial learning: a diffusion MRI study" (from another version of this preprint: version 1, 2025-05-15)

### Supplementary information for “Fornix subdivisions and spatial learning: a diffusion MRI study”

#### S1. Illustrating the cumulative proximity score as a measure of place learning

In our previous report (Hodgetts et al., 2020), we quantified spatial learning using cross-trial changes in subjects’ time to find the hidden target, equivalent to the use of escape latencies in the rodent Morris Water Maze (MWM) (Morris et al., 1982; Vorhees and Williams, 2006). As discussed by Gallagher et al. (2015), latency measures contain no information about the spatial distribution of a subject’s search relative to the escape platform (goal location). This is important because two hypothetical subjects with matched escape latencies could nevertheless differ in the precision of their search; one subject’s search area could be concentrated near the escape platform (low search error), whereas the search area of another subject could be more widely distributed around the MWM (high search error). The performance of these subjects will not be differentiated by latency-based metrics even though the more focused search of the first subject presumably indicates superior place learning.

Gallagher et al. (2015) therefore recommend a more informative trial-wise performance measure, which involves computing the cumulative proximity of the subject to the platform location over the course of the search. With this measure, a subject whose search is concentrated near the platform (low search error) will have a lower final cumulative proximity score compared to a subject with a more widely distributed search trajectory (high search error; see supplementary Figure S1 below). Within a given MWM training trial, cumulative proximity therefore reflects subjects’ search error, and changes in this measure across trials reflects the degree to which subjects improve in their localisation of the target location (Berkowitz et al., 2018).

#### Supplementary Figure S1

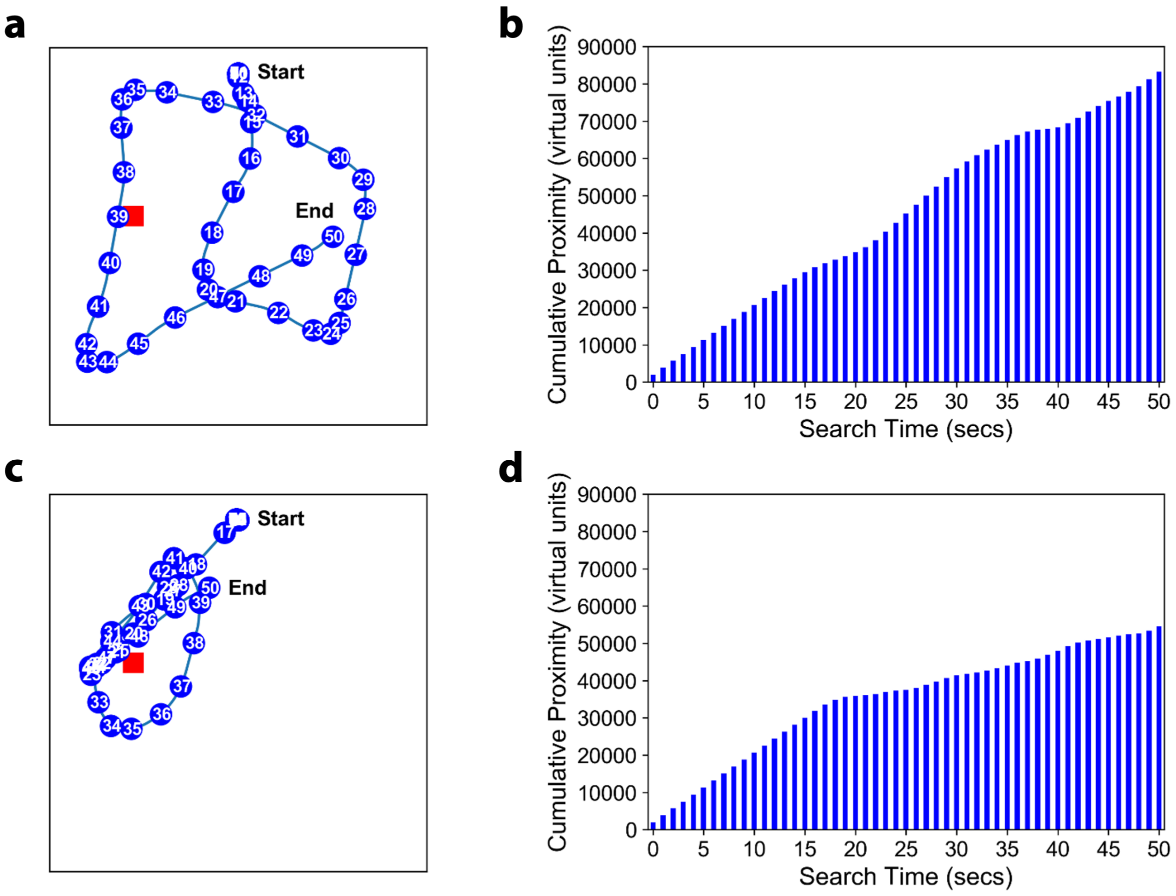


**Figure S1. Representative search trajectories and cumulative proximity during a virtual navigation task based on the Morris water maze.** (A) shows a representative search trajectory (blue line) and the sampling of the participant's proximity to the target location (red square) over 1 second intervals (blue circles). (B) shows the cumulative proximity to the target location over the course of the same trajectory. (C) shows another trajectory which is more precisely focused on the target location, and (D) shows that cumulative proximity accumulates more slowly in this case. For illustrative purposes, these trajectories were truncated from longer trajectories to match the duration over which cumulative proximity was sampled. Neither trajectory therefore intersects the hidden platform. Nevertheless, the final cumulative proximity scores for the trajectories in (A) and (C) were 83,256.6 and 54,524.5, respectively (measured in virtual distance units). This indicates that search trajectories of equivalent duration can be differentiated on the basis of cumulative proximity, with less precise searches characterised by high 'search error' being penalised with a higher cumulative proximity score relative to more spatially precise searches. For a similar example using trajectories matched for total path length rather than search duration/latency, see Gallagher et al. (2015).

**S2: Task learning at the (A) group-level and (B) individual-level. Y-axes represent the cumulative proximity to the hidden sensory. The number of trials (total = 20) is shown on the x-axis.**

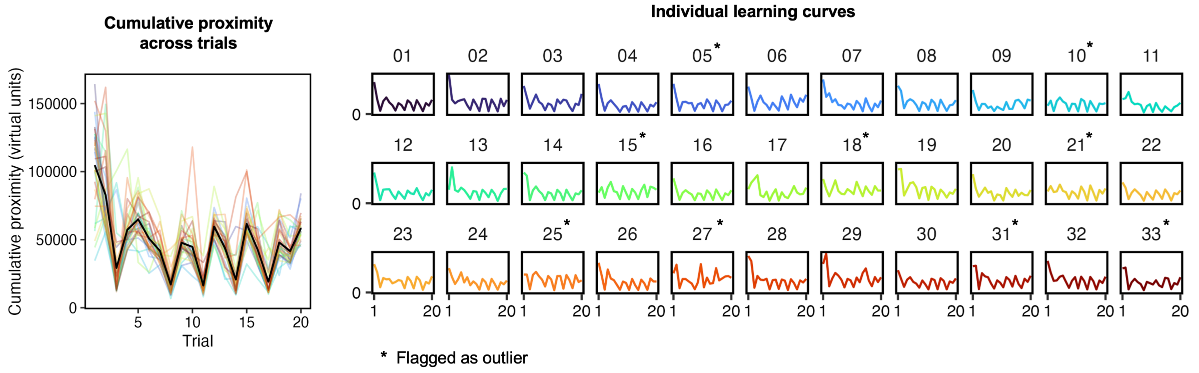


**Figure S2. Navigational learning curves across all trials.** The plot on the left (bold line) displays the average navigational learning curve, illustrating the overall reduction in cumulative search error throughout the task. On the right, individual learning curves for all 33 participants are shown. Participants excluded using our resampling approach are marked with asterisks (see the main text for further details).
